# Assessment of methods for network analysis of single-time-point microbial samples

**DOI:** 10.64898/2025.12.26.696561

**Authors:** Yehonatan Calinsky, Amir Bashan

**Affiliations:** Physics department, Bar-Ilan University, Ramat-Gan, Israel

## Abstract

Harnessing information from a single-time-point microbial sample holds transformative potential for advancing personalized medicine. Traditional techniques, including alpha-diversity (richness-based), beta-diversity (dissimilarity-based) measures and neural network models, effectively pinpoint compositional differences and species abundance variations between the test sample and a reference population. Recently, two novel approaches have been proposed, that, instead of assessing abundance of the individual species, focus on the inter-species relationships in the test sample, the ‘network-impact’ (NI) approach and the individualized dissimilarity-overlap analysis (IDOA). The NI is a measure of the contribution of the test sample to the calculated inter-species correlations in a ‘bottom-up’ approach, while the IDOA analyzes the relationships between species assemblages and their abundances in a ‘top-down’ approach. In this research, we comprehensively evaluate and compare the traditional and the new approaches. Employing the Generalized Lotka-Volterra (GLV) model, we create synthetic samples to rigorously test the classification capabilities of each measure in both supervised and semi-supervised setups. Our findings reveal that when the species of the test sample share similar self-dynamics as the reference cohort but distinct inter-species interactions, they typically can not be classified by conventional dissimilarity-based measures. In contrast, IDOA and NI parameters emerge as successful tools in assessing microbial relationships within a single-time-point sample, while a neural network is highly dependent on the training set size. This study underscores the potential of extracting information from the intricate inter-species relationships based on individual microbial snapshots, paving the way for improving microbiome-based personalized medicine.

## Introduction

The human microbiome, composed of trillions of microbes residing within the body, offers opportunities for advancing personalized medicine [1, 2, 3, 4, 5, 6, 7]. Personalized investigation of an individual’s microbiome can provide critical insights into their unique microbial composition and function, paving the way for more precise and tailored medical treatments.

The abundances of microbial species are determined by a complex network of ecological interactions, including competition for limited resources and cooperative behaviors like cross-feeding [8, 9]. Analyzing this network of inter-species relationships can have significant practical implications. Studies have demonstrated strong associations between certain abnormalities in these interaction networks and a variety of human diseases, such as Crohn’s disease, autism and gestational diabetes [10, 11, 12, 13, 14, 15, 16, 17]. Therefore, analyzing the microbial network associated with the patient’s microbiome and comparing it to the known networks of healthy and diseased populations may provide valuable information about the health status of the patient. Additionally, the effects of actively manipulating the microbiome cannot be accurately predicted without a thorough understanding of the potential changes it may induce in the abundance of other species, which in turn can have a dramatic impact on the entire microbial community.

Analytical methodologies for understanding inter-species relationships, such as correlation analysis or predictive modeling, typically demand a large sample size or prolonged time series with controlled perturbations [18, 19]. However, clinical screenings usually rely on a single-time-point microbial sample due to the time-consuming nature of collecting multiple stool samples. Moreover, given the microbiome’s general stability [7, 20, 21], daily sampling without specific interventions reveals minimal variations that could be primarily noise, hindering our understanding of microbial dynamics.

Thus, the available data in a typical scenario of microbiome-based analysis includes a cohort of samples from different subjects sharing a similar health or disease state (‘reference cohort’), and a single sample from the patient (‘test sample’). Based on these data, the interactions-based diagnosis question is how consistent are the inter-species relationships of the test sample with those associated with the reference cohort.

Traditional methods for comparing a single sample to a reference cohort typically focus on testing the abundance of each species compared to the normal range (represented by the cohort of a healthy population) [22, 23]. The ability of such analysis to detect the effect of inter-species interactions is limited. For instance, as long as the abundance of every species in the test sample aligns within the normal range, it will be classified as ‘normal’, irrespective of the inter-species interactions between the different species.

Recently, two innovative approaches have been introduced to evaluate the consistency of inter-species relationships in test samples with those of reference cohorts: network-impact (NI) parameters [24, 25] and individualized dissimilarity-overlap analysis (IDOA) [26]. While the former adopts a ‘bottom-up’ approach by measuring the test sample’s contribution to the calculated group-level interspecies correlations, the latter employs a ‘top-down’ approach, analyzing relationships between species assemblages and their abundances.

This article systematically analyzes these novel interaction-based methods alongside traditional dissimilarity calculations and neural network models, evaluating their performance using simulated data generated by the Generalized Lotka-Volterra (GLV) model. All methods are rigorously assessed on equal footing to ensure unbiased comparison.

The GLV model describes the dynamics of populations of *n* biological species, including their intra-species and inter-species interactions, as a system of differential equations[27, 28, 29]. Those differential equations are formulated as:

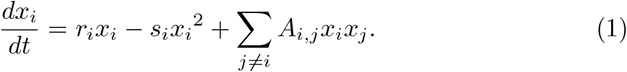

In each equation, *x*_*i*_(*t*) represents the abundance of specie *i* over time. The parameters *r*_*i*_ and *s*_*i*_ denote the self-dynamics, representing the growth rate and the effect of intra-species interactions of that species, respectively. The matrix *A* signifies the interactions among species, where *A*_*ij*_ indicates the effect species *j* has on species *i*.

In this article we use two types of classifications: ‘supervised’ and ‘semi-supervised’. In supervised classification, we construct two (or more) reference cohorts, each comprising of *m* samples which are simulated using the same GLV model, where distinct GLV models are used for each cohort. A test sample is derived from the same GLV model as one of the cohorts. The objective of this setup is to predict the cohort associated with the same GLV model as the test sample, as illustrated in Fig. 1**a**. Different classification methods are compared based on their success rates. This scenario mimics the diagnostic task of classifying individuals into distinct health states.

**Figure 1.**
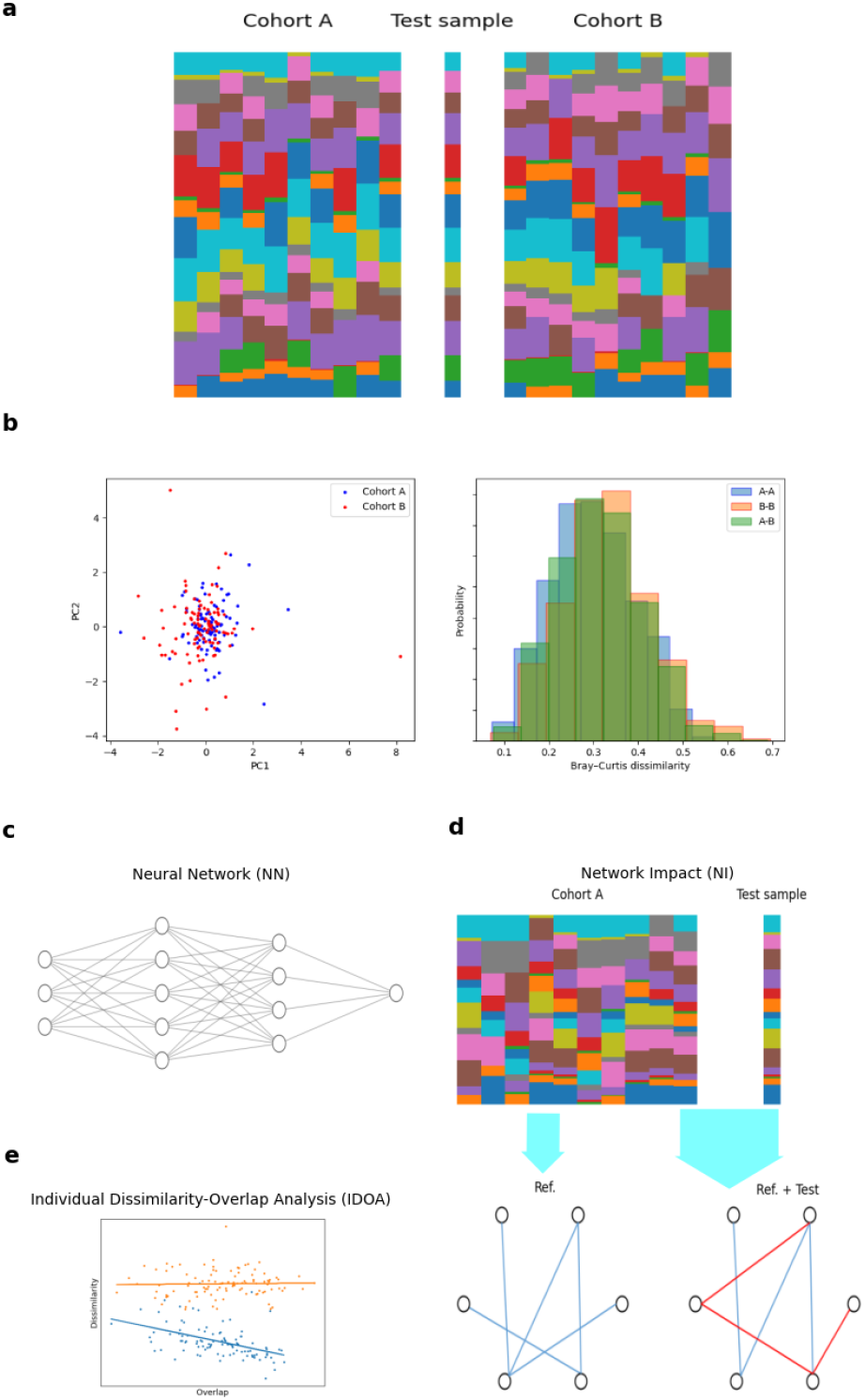
Classification of a single test sample. **a**, An illustration of two ‘reference cohorts’ of samples generated using two different GLV models that share the same self-dynamics (the growth rate *r*_*i*_ and intra-species interactions *s*_*i*_ are the same for species *i* in both models), but with cohort-specific inter-species interactions (off-diagonal elements in *A* matrices). The classification challenge is to determine the original model of a single test sample based on its abundance profile. **b**, The samples of the two reference cohorts are not distinguished using principal component analysis (left panel), and the sample-sample Bray-Curtis dissimilarities between the cohorts are similar to those within the cohorts (right panel). **c**, A neural network model can be trained to classify samples into their original cohorts. The input layer receives the relative abundance vector of the sample and the output layer represents the binary or multi-class classification. **d**, The ‘network impact’ of a single sample is assessed by comparing the correlation networks constructed with and without its inclusion in the reference cohort. In a supervised classification setup, the network impact of the test sample is expected to be lower when calculated with respect to the reference cohort associated with the same GLV model. **e**, Each point represents the dissimilarity-overlap values calculated between the test sample and the samples of a given reference cohort. The IDOA score (the slope of the cloud of points) is expected to be negative when calculated with respect to a cohort associated with the same GLV model (blue) and zero when calculated with respect to an unrelated cohort (orange).

In the semi-supervised classification, only one cohort is utilized, and the test sample can either be generated from the same GLV model or be a ‘shuffled sample’ based on this cohort (as explained in the Methods section). Each classification method is then evaluated using ROC curve to indicate its ability to distinguish between real and shuffled samples. This setup represents a scenario of regular health screenings, where a single-time-point microbial sample is compared solely to a cohort of healthy samples.

## Methodology

In this article, we use different types of classification approaches.

### Dissimilarity-based methods

For each dissimilarity measure employed, we compute the dissimilarity between the test sample and each sample in the cohort. Specifically, we used Euclidean distance and Bray-Curtis dissimilarity. The predicted cohort is determined by the cohort that yields the minimal mean dissimilarity with the test sample. This process is carried out separately for each dissimilarity measure.

### Neural Network (NN) model

In the Neural Network method, we establish a neural network, which is then trained using the different reference cohorts to subsequently predict the class (the associated reference cohort) of the test sample (see Methods section for the detailed description of the network architecture). The output nodes of the neural network represent the probabilities of the test sample being associated with each reference cohort. The predicted cohort is determined by the cohort associated with the highest probability.

### IDOA method

The IDOA method is originally introduced in Ref. [26]. In this method, the IDOA value is determined by calculating the overlap and dissimilarity between the test sample and each of the samples in a reference cohort and taking the slope of a linear regression model that represents the relationships between dissimilarity and overlap for all sample pairs with overlap larger than 0.5.

When the test sample originates from a different GLV model than the reference cohort, the dissimilarity and overlap are expected to be independent, resulting in a slope close to zero. In the case of a shuffled sample, the dissimilarity and overlap should also be uncorrelated, resulting in a near-zero slope. Conversely, if the test sample is generated from the same GLV model as the reference cohort, the dissimilarity and overlap should demonstrate dependence, yielding a negative slope indicating greater overlap with lower dissimilarity.

### Network Impact (NI) parameters

The ‘network impact’ of a single sample is assessed by comparing the correlation networks constructed with and without its inclusion in the reference cohort. The difference between the two correlation networks is then evaluated using three distinct parameters: structural difference (NI-SD)[24], weighted difference (NI-WD), and ‘theta’ (NI-T)[25]. In this manuscript, in addition to the original definitions used in Ref. [25], we introduced also new versions for the weighted difference and the theta parameters (see Methods).

The rationale behind the NI method is analogous to ‘drop-one-out’ experiments, where the exclusion of a data point that is quite similar to the group does not affect much its averaged properties. Similarly, when the test sample was generated using the same GLV model as the rest of the samples in the reference cohort, its exclusion is not anticipated to change much the correlation network. However, a noticeable difference is expected to the correlation network when the test sample was generated using different GLV model.

## Results

In this section, we present the evaluation results of the methods in the context of supervised and semi-supervised classification tasks.

### Supervised classification

The supervised classification task involves assigning a single test sample into one of several reference cohorts generated using distinct GLV models. Each reference cohort comprises *m* simulated ‘samples’, with the GLV models sharing the same set of *r*_*i*_ and *s*_*i*_ values to maintain consistency of characteristic species abundances across the different reference cohorts. However, unique interaction matrices (*A*_*i,j*_ for *i* ≠ *j*) are selected for each model using the same density of interactions and interaction strength (See Methods). Note that the abundance profiles of the different reference cohorts are similar as long as the inter-species interactions are not too strong, as shown for the binary case in the Methods section (Fig. 5). The objective is to identify the associated cohort based solely on the abundance profiles of the test sample and of the reference cohorts, while the underlying GLV models are assumed unknown. For all methods except the NN, the classification is performed in two steps: first, a score is calculated independently for the test sample with respect to each of the reference cohorts. Then, the test sample is classified into the cohort associated with the minimal mean distance, minimal network-impact or most negative IDOA, respectively. In contrast, the NN is firstly trained using all the reference cohorts, and then can provide the probabilities of the test sample belonging to each cohort.

### Binary Classification

Our evaluation begins with binary classification, where two cohorts of equal size (*m*) are generated along with a test sample. In each classification test, each method is provided with two reference cohorts and a single test sample and its output is the predicted reference cohort associated with the test sample. This step is repeated 100 times with different test samples, and the success rate is defined as the proportion of correctly classified test samples. In an ideal case, the reference cohorts contain large number of samples, where all the samples within the same cohort follow the same ecological dynamics (i.e., ‘universal dynamics’ [30]). In real-world scenarios, however, only finite reference cohorts are available, and the underlying dynamics may vary across different individual samples within the same cohort.

In this section, We study the effect of these two basic properties: the number of samples *m* in the reference cohorts and the level of sample-to-sample variability of the underlying dynamics (‘biological noise’). The later property is represented by introducing sample-specific stochasticity (a noise value *δ*, see Methods) to the equations representing an individual sample.

First, we study the effect of the size of the reference cohorts *m* on the success rates of the different methods (the value of *δ* was randomly sampled from a uniform distribution ranging between 0 and 0.3). As shown in Fig. 2**a**, the dissimilarity-based methods perform the worst. The Euclidean method achieves a success rate of approximately 50%, independent of *m*, which is no better than random guessing. The Bray-Curtis method performs better, with success rate that improves from around 50% to 70% as *m* increases to 500. The NI-T1 and NI-T2 methods show success rates of roughly 75% and 60%, respectively, with minimal dependence on *m*. The NI-SD method achieves an approximately 88% success rate for *m* between 25 and 90, but this declines for larger *m*, indicating an optimal reference cohort size for this method. These results align with those reported in Ref. [25] for the NI-T and NI-SD parameters. The success rate of the NN method gradually increases with *m*, reaching around 82% when *m* = 500. Notably, the IDOA method and both variations of the NI-WD significantly outperform the other approaches, with success rates of approximately 96%-97% for *m* = 25, maintaining high accuracy as *m* grows larger.

**Figure 2.**
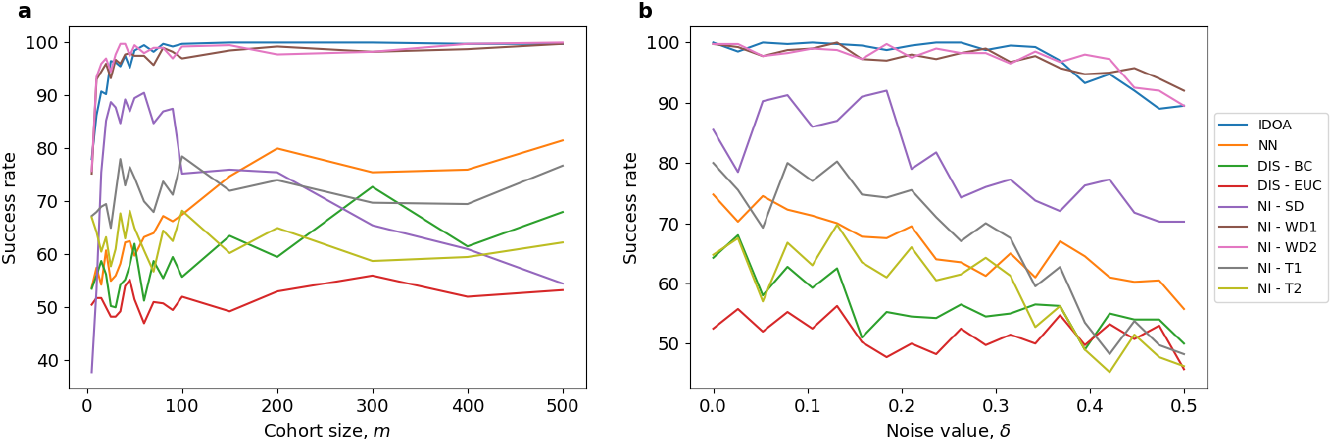
Success rates of methods for binary supervised classification. Two reference cohorts, each with *m* samples of *n* = 100 species, were generated using cohort-specific GLV models. 100 test samples were generated, where each was generated using one of the two GLV models. The success rate is defined as the percentage of correctly classified test samples into the reference cohort associated with the same GLV model. Finally we average the success rates over 4 independent realizations. **a**, The success rate versus the size of the cohorts, *m*. The IDOA, NI-WD1 and NI-WD2 exhibit high success rates even for small size of reference cohort compared with the other methods. **b**, The success rate versus the noise value, *δ*, i.e., sample-to-sample variability in the GLV parameters within the same cohort. Here, *m* = 100 samples in each cohort. The IDOA, NI-WD1 and NI-WD2 exhibit higher performances even in the presence of biological noise in the reference cohorts compared with the other methods.

These findings indicate that the binary classification setup, which is designed to emphasize inter-species dynamic variations, is best addressed by interaction-based methods such as IDOA and the weight-difference network impact (NI-WD1 and NI-WD2). These methods require relatively small number of reference cohorts to achieve high success rates.

Next, we evaluate how different methods respond to different levels of noise *δ* (with fixed cohort size *m* = 100), as shown in Fig. 2**b**. Generally, as the noise level increases, the success rate of all methods declines. Yet, while for most methods the success rate declines even for relatively low level of noise, the success rates of the IDOA and the weight difference variations of Network Impact (WD1 and WD2) remain high up to *δ* = 0.3. This relatively higher noise-resilience of IDOA and both NI-WD variations may be attributed to their focus on inter-species interactions, which is the primary discrimination of the two cohorts in this setup.

### Multi-Class Classification

Here, we explored the effectiveness of the methods with *k* (2 *≤k ≤*5) reference cohorts, each composed of *m* = 100 samples. In each classification test, each method is provided with *k* reference cohorts, simulated using distinct GLV models, and a single test sample simulated using one of the models. The output of each classification method is the predicted reference cohort associated with the test sample, and the success rate is averaged over 100 independent test samples. Similar to the binary classification setup, the different GLV models share identical self-dynamics for each species, therefore the reference samples have similar characteristic abundance profiles, as demonstrated in the PCA plots in Fig. 3**a-d**.

**Figure 3.**
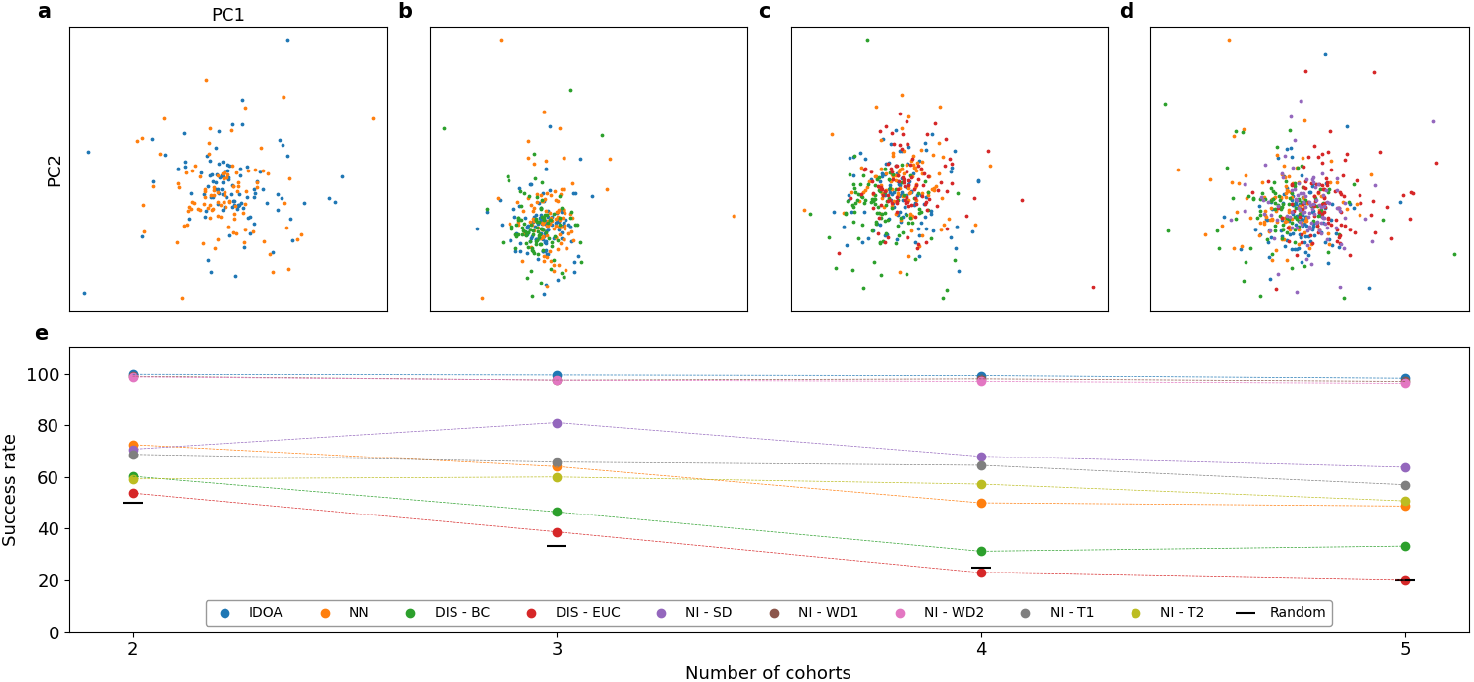
Comparison of methods in multi-class supervised classification setup. Each cohort consists of *m* = 100 samples of *n* = 100 species. The GLV models of all the cohorts share the same self-dynamics but differ in their inter-species interactions. **a-d**, PCA plots of the reference samples colored according to their associated cohorts. In all cases, the clouds of the different cohorts largely overlap, i.e., a single test sample cannot be classified into its underlying GLV model using its sample-sample similarity to the reference cohorts. **e**, The success rate of each method versus the number of reference cohorts, averaged over 4 different realizations. The black lines represent the expected success rate of random guesses (100*/k* %).

The success rates for each classification method, as a function of *k*, are presented in Fig. 3**e** for 2 *≤k ≤*5. The methods are compared with the expected success rate under random guessing (100*/k* %), marked with black line for the different *k* values. The success rates of the dissimilarity-based (DIS-BC and DIS-EUC) and the NN methods tend to decline with *k*, while the interaction-based methods (IDOA and all NI parameters) displayed less sensitivity to the number of cohorts.

### Semi-supervised classification

In this section, we explore the added value of the inter-species interactions in the case where only a single reference cohort is available. To focus on the inter-species interactions, we generate two types of samples: 1) ‘real samples’ - samples simulated using the same GLV model as the reference cohort, and 2) ‘shuffled samples’, generated by shuffling the abundances of the reference samples, as detailed in the Methods section. The shuffled samples preserve the characteristic species abundances of the real samples, but lack any inter-species interrelations (as demonstrated in Fig.4**a**). Therefore, distinguishing between GLV samples and shuffled samples highlights the contribution of inter-species interactions to the analysis of single-time-point samples. The semi-supervised setup represents a scenario where the health state of a subject is assessed based on a single-time-point sample with respect to a single reference cohort of healthy subjects.

In this setup, a single score is calculated for each test sample with respect to the reference cohort, composed of *m* = 100 samples simulated using a specific GLV model. Here, we generate 100 test samples, each of them is either real or shuffled, and the methods’ ability to distinguish between real and shuffled samples is evaluated in terms of the receiver operating characteristic (ROC) curves. For the NN, the model is trained using the reference cohort as well as a separate ‘shuffled cohort’ consisting of *m* = 100 shuffled samples, and the score is the probability for the class associated with the original reference cohort.

Figure 4**b** illustrates the distribution of the classification scores obtained from each method for real samples and for the shuffled samples. IDOA and NI-WD2 exhibit a clearer distinction between the distributions of scores calculated for the real and shuffled samples compared with other methods, for which they largely overlap. However, the scores of IDOA and NI-WD2 calculated to the same individual samples are not correlated, where many of the real samples are distinguished by only one of the two methods, as shown in Fig. 4**c**. This result suggests that the ‘top-down’ (IDOA) and the ‘bottom-up’ (NI-WD2) interaction-based approaches capture different aspects of the inter-species interrelations in real samples.

**Figure 4.**
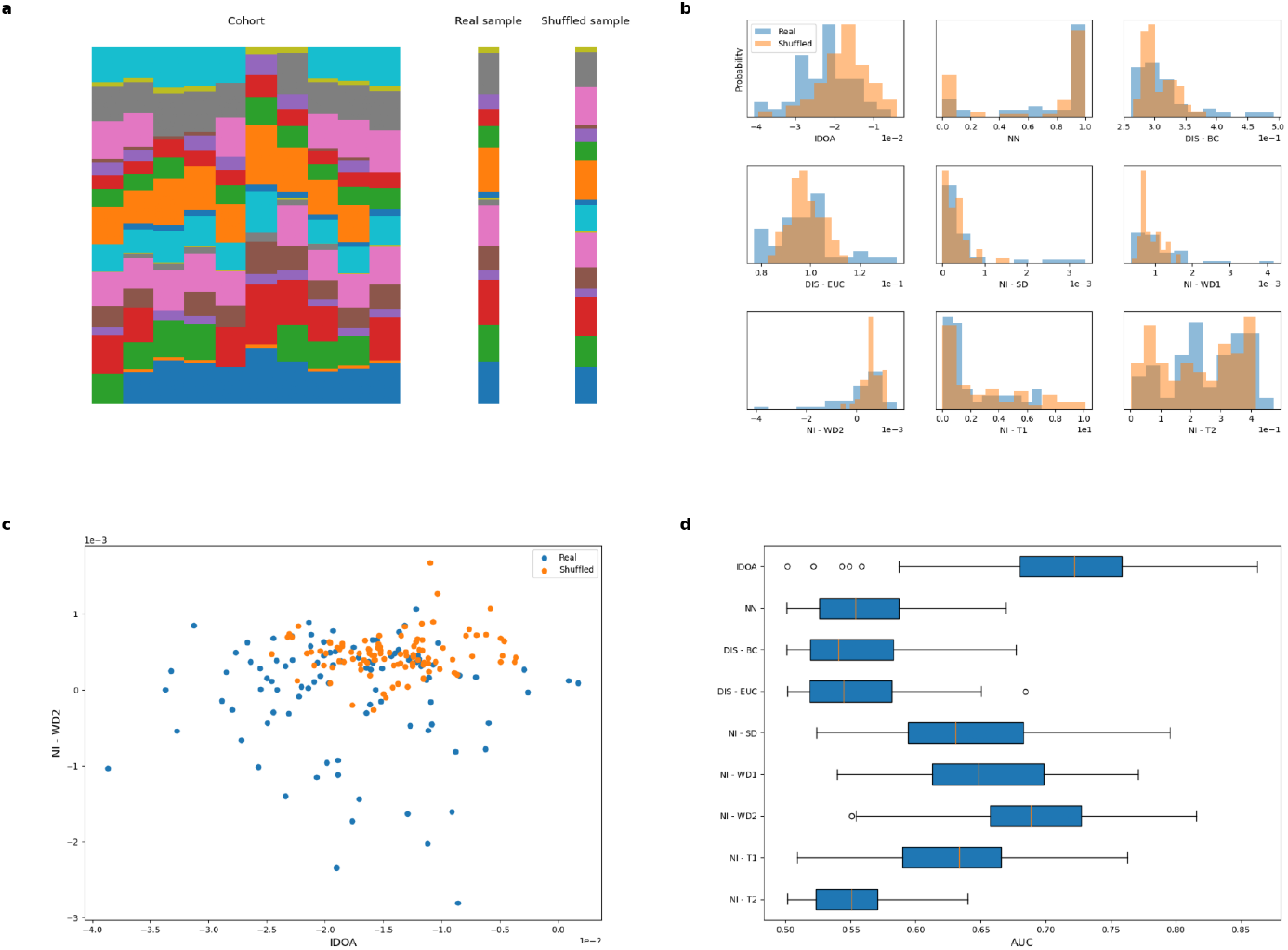
Semi-Supervised Classification of a single microbial sample. ‘Real’ samples are generated using the same GLV model as the reference cohort and ‘shuffled’ samples are generated by shuffling the abundances of the reference samples (See Methods). **a**, A schematic example of the semi-supervised classification setup in a system of *n* = 20 species. Each vertical stack bar represents the relative abundance profile of a single sample. The species abundances in the shuffled sample mimic those of the reference samples but with no inter-species interactions. **b**, Distributions of the output values of the different methods calculated for 100 test samples with respect to a reference cohort consisting of *m* = 100 samples with *n* = 100 species. Every test sample is randomly generated as either ‘real’ or ‘shuffled’. **c**, Each point represents the IDOA and NI-WD2 values calculated for the same single test sample. While the IDOA and NI-WD2 values calculated for about half of the ‘real’ test samples are clearly smaller than the ‘shuffled’ samples, the scores seem to be uncorrelated suggesting that the ‘top-down’ and ‘bottom-up’ approaches capture different aspects of the inter-species relationships. **d**, Area-under-curve (AUC) values of the Receiver Operating Characteristic (ROC) curves associated with each of the methods, calculated for 100 realizations. Box plots represent the first and third quartiles; middle line, median; circles, outliers.

Figure 4**d** presents the Area Under the Curve (AUC) values of each methods for 100 realizations. The AUC values of the IDOA method are markedly higher compared with the other methods, the Network Impact parameters have intermediate AUC values, and the dissimilarity-based methods and neural network model have the lowest AUC values. The NN method’s suboptimal performance in the semi-supervised classification task, compared to the supervised classification, may be due to its reliance on well-defined class boundaries, which are absent in the case of shuffled samples.

## Discussion

In the realm of personalized medicine, where rapid and accurate diagnoses are essential, there is an increasing demand for accessible analysis tools capable of handling single-time-point microbial samples. This study evaluates the performance of various methods for analyzing inter-species interactions within microbial communities. Our tests are motivated by two central types of real-world diagnostic scenarios, with supervised classification mimicking diagnostic task with multiple health states or diseases and semi-supervised classification analogous to regular health screening.

This manuscript presents two primary findings: first, it systematically compares the performance of different methods in analyzing single-time-point samples with an emphasis on interactions-based approaches. Second, it contrasts two analytical paradigms for complex systems: the ‘bottom-up’ and ‘top-down’ methods.

We used different evaluation metrics for each classification setup. In the supervised setup, the classification of a test sample depends entirely on comparisons of scores calculated for a test sample with respect to each reference cohort, with performance measured by success rates. However, in the semisupervised setup, where only a single score is calculated per test sample, classification potential is assessed through ROC curves and the corresponding AUC values.

Our results show that both the IDOA and NI-WD2 methods outperform other approaches in both classification setups. These results demonstrate that the dedicated interactions-based methods can capture inter-species relationships that other methods, which focus on individual species abundances, tend to overlook.

Specifically, in the supervised setup, both IDOA and NI-WD2 yield high success rates, making each of them a reliable measure, independently. However, in the semi-supervised context, they seem to capture different aspects of the data. Qualitatively, while the IDOA measures the consistency of the dynamics of a test sample with respect to a reference cohort, the NI method assesses the magnitude of their differences. This suggests that integrating the information from both methods could yield optimal results.

We also evaluated the performance of a neural network (NN) in the classification tasks. Although we chose specific model characteristics (e.g., architecture, depth), which may influence the quantitative outcomes, some qualitative insights can be drawn. In the supervised setup, the NN shows moderate success, with performance improving as the size of the reference cohorts increases. However, it seems that much larger training sets would be required to match the success rates of the dedicated interactions-based methods. Another key limitation of the NN is its lack of interpretability; it does not reveal whether its classifications are driven by inter-species interactions or subtle abundance variations in specific species. In the semi-supervised setup, the NN’s performance was considerably weaker due to the absence of well-defined reference cohorts, further highlighting its limitations in this context.

In this study, both reference data and test samples were generated to focus on inter-species interactions by keeping the self-dynamics of individual species constant. However, in some real-world scenarios, single-time-point microbial samples may be classified based on simple similarity measures rather than inter-species interactions. In practice, we recommend first examining whether reference cohorts have distinguishable characteristic abundance profiles. If so, similarity-based measures may suffice, but interaction-based methods could provide complementary insights to improve classification accuracy and can be integrated with the other methods.

The two leading interaction-based methods, IDOA and NI, represent distinct approaches to analyzing complex systems. NI follows a ‘bottom-up’ strategy, relying on detailed correlation networks to assess test samples. This approach may provide additional insights beyond the required for the classification setup, such as characterization of the interrelations of individual species and assessment of the global network structure. However, the computational complexity of this approach can be significant, as it requires extensive calculations. Therefore, the choice between a ‘bottom-up’ approach, such as NI, and a ‘top-down’ approach, like IDOA, will depend on the specific depth of analysis required and the computational resources available. While NI may offer richer information, IDOA provides a more streamlined, faster method for classification tasks where detailed network analysis is unnecessary.

Our results do not aim to present definitive conclusions, but rather qualitative insights about the different methods, since each of the assessed methods allows for numerous technical variations. For example, future work could explore additional dissimilarity measures within the IDOA framework or test alternative neural network architectures and activation functions. Furthermore, the specific setup designed here could be generalized by experimenting with different shuffling techniques for semi-supervised classification, and the results for samples simulated using the GLV model should be tested on non-GLV models, such as Consumer-Resource models and real-world data.

The primary contribution of this study is to establish a systematic framework for comparing these methods and illustrating their potential for analyzing single-time-point microbial samples. This framework paves the way for further advancements in personalized medicine diagnostics, particularly in understanding complex microbial interactions.

## Methods

We begin by outlining the process for simulating samples using the Generalized Lotka-Volterra (GLV) model, followed by a method for generating shuffled samples. Afterward, we describe the four main approaches for evaluating individual samples: IDOA, NI, NN, and dissimilarity-based methods. Also, the computational complexity of each method is analyzed based on the number of species *n* and the number of samples *m* in a reference cohort.

### GLV simulations

We define a ‘sample’ as the steady state of a particular GLV model described in Eq. 1, where each sample is represented as a R^*n*^ vector, such that ***x*** = (*x*_1_, …, *x*_*n*_) *∈* R^*n*^. In our simulation, the intrinsic growth rate of species *i* is randomly chosen from a uniform distribution *r*_*i*_ *∈* 𝒰 (0, 1). We also set *∀i ∈* [1, *n*] : *s*_*i*_ = 1. As for the matrix *A*, the off-diagonal elements are drawn from a uniform distribution *A*_*i,j*_ *∈* 𝒰 ( *− β, β*) with probability *C* and zero otherwise. The default parameters in this article are *C* = 0.1 and *β* = 0.025 (unless otherwise specified). As illustrated in Fig 5, the cohorts generated with these parameters exhibit comparable abundance profiles, even though their interaction structures differ significantly.

**Figure 5.**
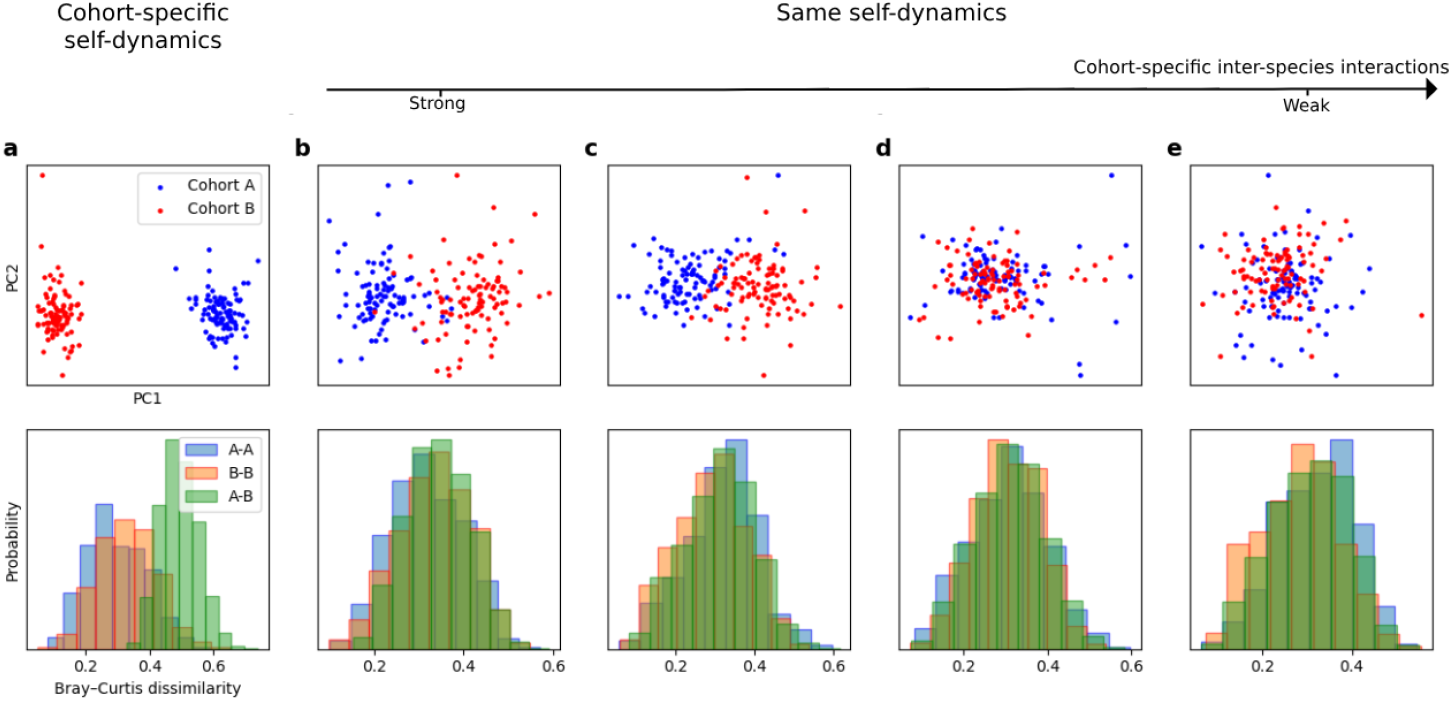
Creating reference cohorts with shared characteristic abundances but varied inter-species interactions. Two cohorts, each composed of *m* = 100 samples, are generated using two different GLV models. Top row: Each point represents the abundance profile of a single sample on the plane of the two leading principal components, colored according to its associated cohort. Bottom row: Distributions of Bray-Curtis dissimilarity calculated between samples of cohort *A* (blue), between samples of cohort *B* (orange), and between samples from different cohorts (green). **a**, When the GLV models have distinct self-dynamics, the inter-cohort dissimilarity is significantly larger than the intracohort dissimilarity. Thus, a single sample can be effectively identified by its similarity to the cohorts. **b-e**, The two cohorts share the same self-dynamics (the growth rate *r*_*i*_ and intra-species interactions *s*_*i*_ are the same for species *i* in both models) but cohort-specific inter-species interactions (*β* = 0.1, 0.04, 0.025, 0.015, respectively). When the inter-species interactions are weak, the separation of the two cohorts by the sample-sample similarities and PCA is practically unnoticeable.

To simulate a cohort of samples, we first define ‘base parameters’, a set of *r*_*i*_ for all *i* and matrix *A*^*∗*^. Then, we simulate an individual sample *ν* in that cohort, using parameters that are variations of the base parameters with random noise parameterized by *δ*. Specifically, we define *A*^*ν*^ = (1 *− δ*^*ν*^) *· A*^*∗*^ + *δ*^*ν*^ *· B*^*ν*^, where *B*^*ν*^ is a sample-specific matrix randomly generated for each sample using the the same *C* and *β* parameters as *A*^*∗*^ and *δ*^*ν*^ is randomly selected from the distribution 𝒰 (0, *δ*_*max*_). Unless specified differently, we use *δ*_*max*_ = 0.3 (In Fig. 2**b** we systematically study other noise strengths).

To generate each simulated sample, we randomly initialize a species assemblage with abundances drawn from a uniform distribution. The average prevalence for each sample *p*^*ν*^ is set to a value randomly selected from the uniform distribution 𝒰 (0.6, 0.9). Individual species are assigned zero abundance with a probability 1 *− p*^*ν*^; otherwise, their abundances were sampled uniformly from 𝒰 (0, 1). We then integrate Eq.1 using the *odeint* function from *scipy*.*integrate* library of Python. We repeat the above steps to generate *m* samples, each represents the steady state abundance of *n* species. The cohort is represented by *X ∈* R^*m×n*^ matrix.

At last, we normalize the steady states in order to represent a relative abundance samples, such that 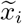 represents the relative abundance of species *i* and

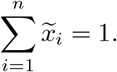

Finally, the generated cohort is represented by 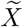 normalized samples. matrix that contains the

### Generating shuffled samples

We create an individual shuffled sample by taking the non-normalized reference cohort *X*. The shuffled sample ***y***^*ν*^ *∈* R^*n*^ is defined by: 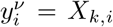 while *k* is randomly selected from [1, *m*] with equal probability. This process preserves the characteristic abundance of each species but removes any correlations between the species. We then normalize the shuffled sample. We repeat these steps to generate the required number of shuffled samples.

### Individual Dissimilarity-Overlap Analysis (IDOA)

IDOA calculates the similarity between a test sample and the reference cohort based on the Dissimilarity-Overlap Curve (DOC). For each pair of samples, the overlap quantifies the similarity of shared species *S* = *{i* | *x*_*i*_ *>* 0 and *y*_*i*_ *>* 0*}*:

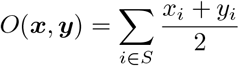

The dissimilarity is computed using the Jensen-Shannon Divergence on the re-normalized samples with respect to the shared species 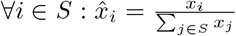 and 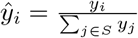. The dissimilarity value is then defined by:

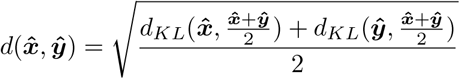

and 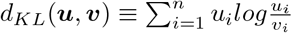.

#### Computational Complexity

Here, we compute the dissimilarity-overlap points for the test sample with each sample in the reference cohort. Initially, we determine the subset of all species existing in both the test sample and the cohort sample, requiring *O*(*n*) time. Subsequently, each dissimilarity-overlap pair calculation takes *O*(*n*) time. This results in a total complexity of *O*(*n·m*) for the DOC curve calculation. Finally, polynomial fitting is applied to determine the IDOA value, an operation that is negligible based on our simulations in all tested values of *n* and *m*. Altogether, the complexity is approximately *O*(*n · m*) (neglecting the linear regression task).

### Network Impact (NI)

To construct a correlation network for a given cohort, represented by a *N ∈* R^*n×n*^ matrix, we define *ρ*_*i,j*_ as the Pearson correlation coefficient of species *i* and *j*, considering only the subset of samples in the cohort where both species exhibit non-zero abundance. The matrix element *N*_*i,j*_ is assigned the value of *ρ*_*i,j*_ if the corresponding p-value is less than 10^*−*3^; otherwise, it is set to 0.

Next, we define the unweighted correlation network *N* ^*′*^, described by the adjacency matrix, as 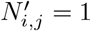 if *N*_*i,j*_ ≠ 0, and 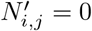 otherwise.

Let *N* ^(*m*)^ represent the correlation matrix of the reference cohort, and *N* ^(*m*+1)^ represent the correlation matrix of the combined set of the reference cohort and the test sample. Similarly, *N* ^*′*(*m*)^ and *N* ^*′*(*m*+1)^ denote the unweighted correlation networks for the reference cohort and the combined cohort plus test sample, respectively. The shared links between the correlation networks are denoted as the set 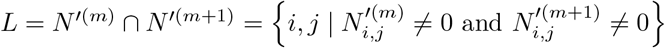.

We quantify the differences between *N* ^(*m*+1)^ and *N* ^(*m*)^ using five parameters, where NI-SD, NI-WD2 and NI-T2 were introduced in Ref. [25]. These parameters are categorized into three types:

#### Structural Difference

First, we assess how much the structure of the unweighted network *N* ^*′*(*m*+1)^ deviates from *N* ^*′*(*m*)^ using the NI-SD parameter. This difference is captured using the Jaccard dissimilarity between the two unweighted networks:

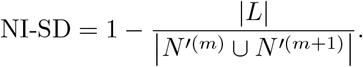

#### Weight Difference

Second, we focus on the shared links and evaluate, using the NI-WD parameters (NI-WD1 and NI-WD2), the degree of variation in their weights. Two variations are defined to measure the differences between the weights of the shared links:

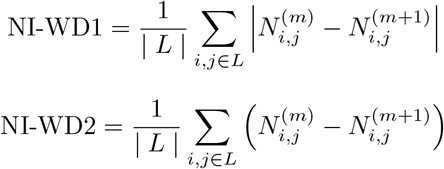

NI-WD1 equals zero only when all the links weights are equal in *N* ^(*m*)^ and *N* ^(*m*+1)^, while NI-WD2 equals zero when the overall increased and decreased links’ weights between the two networks cancel each other.

#### Theta

Lastly, NI-T parameters (NI-T1 and NI-T2) quantifies the proportion of network links whose weights decreased after the test sample was added to the reference cohort. For that, we define the subset of links that change between the networks as

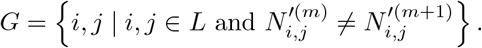

The subset of links that exhibit a decrease in weight is defined as

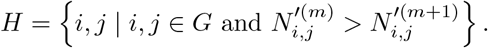

Two variations of the *θ* parameter are introduced:

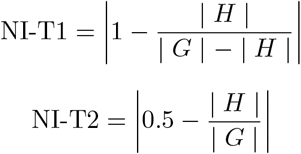

While both NI-T1 and NI-T2 score the ‘neutral’ case where 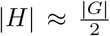 is zero, the difference between them is that NI-T2 is linear and symmetric with respect to 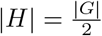 and NI-T1 is non linear and gives higher score for the case of 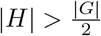.

#### Computational Complexity

To compute each network impact parameter, we first calculate the interaction network of each cohort (or cohort + test sample). In the interaction network, we compute the Pearson correlation for each pair of species. The Pearson correlation calculation requires Ω(*n* + *m*) time, hence the entire network calculation takes Ω(*m · n*^2^ + *n*^3^) time. For each test sample, we compute the interaction network with each cohort. The complexity here represents a lower bound since Python does not precisely reflect the exact calculations made, but this lower bound is necessary. Subsequently, each parameter (Structural Difference, Weight Difference, and Theta) requires about *O*(*n*^2^) time for comparing the networks with and without the test sample. Overall, the computational complexity for each parameter is Ω(*m · n*^2^ + *n*^3^).

### Neural Network (NN)

The neural network utilized in this study comprises 2 hidden layers, with 50 and 100 neurons, respectively. It’s worth noting that altering the number of layers and neurons does not impact the results, provided the hidden layers contain a minimum of 50 neurons (for the simulated data). We employ the *Keras* module in Python for configuring, training, and employing the Neural Network.

The output of the neural network for each class (reference cohort) is a value ranging from 0 to 1. A higher value signifies that the network predicts the test sample is more likely to belong to this class. For supervised classification, we designate the cohort with the highest value as our prediction. In semi-supervised classification, we generate a ‘shuffled cohort’ consisting solely of shuffled samples. Subsequently, we train the network using both the real and shuffled cohorts. The calculated value represents the output for the real cohort, indicating the network’s level of certainty regarding whether the sample is associated with the real cohort.

#### Computational Complexity

The complexity of neural networks varies based on the specific network architecture and the specific training process. Generally, while the classification time is brief, the training time is significantly longer.

**Table 1:** Summary of all methods.

| Abbreviation | Method Name | Complexity |
| --- | --- | --- |
| IDOA | Individual Dissimilarity-Overlap Analysis | $O(n \cdot m)$ |
| NN | Neural Network | - |
| DIS - BC | Bray-Curtis Dissimilarity | $O(n \cdot m)$ |
| DIS - EUC | Euclidean Distance | $O(n \cdot m)$ |
| NI - SD | Network Impact - Structural Difference | $\Omega(m \cdot n^2 + n^3)$ |
| NI - WD1 | Network Impact - Weight Difference 1 | $\Omega(m \cdot n^2 + n^3)$ |
| NI - WD2 | Network Impact - Weight Difference 2 | $\Omega(m \cdot n^2 + n^3)$ |
| NI - T1 | Network Impact - Theta 1 | $\Omega(m \cdot n^2 + n^3)$ |
| NI - T2 | Network Impact - Theta 2 | $\Omega(m \cdot n^2 + n^3)$ |

### Dissimilarity-based measures

Here we utilize two dissimilarity measures for comparing a sample to a cohort: Euclidean distance and Bray-Curtis dissimilarity.

#### Euclidean Distance

This metric calculates the ‘straight-line’ distance between two samples in an n-dimensional space. It is computed using the formula:

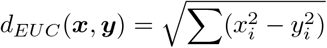

#### Bray-Curtis Dissimilarity

This dissimilarity evaluates the proportion of species that differ in abundance between two samples. It is computed using the formula:

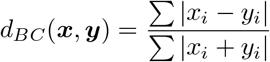

#### Computational Complexity

Both dissimilarity methods require *O*(*n · m*) time to calculate the dissimilarity of the test sample with each sample in the cohort. Where the dissimilarity is averaged over a cohort of *m* samples, an additional *O*(*m*) time is needed to compute the mean distance.

### Source Files

All the Python files in this research can be found in this project:

https://github.com/yonikal56/single-sample-analysis-comparison.

